# Manganese complex [Mn(CO)_3_(tpa-κ^3^*N*)]Br increases antibiotic sensitivity in multidrug resistant *Streptococcus pneumoniae*

**DOI:** 10.1101/2020.02.27.969048

**Authors:** Apostolos Liakopoulos, Roberto M. La Ragione, Christoph Nagel, Ulrich Schatzschneider, Daniel E. Rozen, Jonathan W. Betts

## Abstract

The emergence of multidrug-resistance (MDR) in *Streptococcus pneumoniae* clones and non-vaccine serotypes is of increasing concern, necessitating the development of novel treatment strategies. Here, we determined the efficacy of the Mn complex [Mn(CO)_3_(tpa-κ^3^N)]Br against MDR *S. pneumoniae* strains. Our data showed that [Mn(CO)_3_(tpa-κ^3^*N*)]Br has *in vitro* and *in vivo* antibacterial activity and has the potential to be used in combination with currently available antibiotics to increase their effectiveness against MDR *S. pneumoniae*.

## Introduction

*Streptococcus pneumoniae* is the main cause of community-acquired pneumonia, meningitis and bacteremia in children and adults (1), with pneumonia remaining the leading cause of death in children under 5 years of age worldwide (2). In addition, pneumococcal disease causes the most deaths among vaccine-preventable diseases, according to the World Health Organization (WHO) (3). Although the available pneumococcal vaccines have reduced invasive pneumococcal disease (IPD), current vaccines only protect only against a fraction of the more than 97 circulating serotypes. Eradication of the vaccine-included serotypes has caused rapid serotype replacement, followed by an increase in the carriage, prevalence and disease caused by non-vaccine serotypes (4). As a consequence of the incomplete protection against circulating serotypes, antibiotic therapy remains a mainstay of IPD treatment (5).

The emergence of multidrug-resistant *S. pneumoniae* strains worldwide compromises the available treatment options for IPD (6–11) and imposes the need for alternatives to traditional anti-pneumococcal agents. Managanese-carbonyl complexes, such [Mn(CO)3(tpa-κ^3^*N*)]Br, have been proposed as antibacterials against Gram-negative bacteria (12, 13), especially in combination with membrane permeabilisers like colistin (14). Although their mechanism of action is still elusive, a combination of membrane disruption, interference of metal ion uptake and inhibition of respiration has been proposed (15). Previous data suggests that the manganese-coligand core of the title compound does not reach the intracellular environment in Gram-negative bacteria, possibly due to the inability of the compound to cross the outer bacterial membrane (16), but their antibacterial activity against Gram-positive bacteria is yet to be explored.

The aim of this study was to evaluate the *in vitro* and *in vivo* activity of manganese complex [Mn(CO)_3_(tpa-κ^3^*N*)]Br alone or in combination with commonly used antibiotics against multidrug-resistance strains of *S. pneumoniae*.

## Material and Methods

### Bacterial Strains, growth conditions and media

A total of 20 human-derived clinical non-duplicate invasive and multidrug resistant *S. pneumoniae* strains were provided by the CDC *Streptococcus* Laboratory and were included in the study. Their relevant characteristics are indicated in Table S1. *S. pneumoniae* strains were grown in cation-adjusted Mueller-Hinton broth (BD, New Jersey, USA) supplemented with 100 U of catalase (Worthington Biochemical Corporation, New Jersey, USA) and 20 mg/L β-NAD (Sigma-Aldrich, St. Louis, USA) at 37°C under 5% CO_2_ for 18-24 h. Blood agar plates were made from Tryptic soy agar (BD, New Jersey, USA) with the addition of 0.5% yeast extract (BD, New Jersey, USA) and 5% defibrinated horse blood (Sanbio, Uden, The Netherlands). [Mn(CO)_3_(tpa-κ^3^N)]Br (USC-CN028) was synthesised according to a previously published procedure (13).

### MIC and MBC determination

Minimum inhibitory concentrations (MICs) were determined in triplicate by broth microdilution according to European Committee on Antimicrobial Susceptibility Testing (EUCAST; http://www.eucast.org) and ISO 20776-1:2006 guidelines with the exception that cation-adjusted Mueller-Hinton broth (BD, New Jersey, USA) was supplemented with 100 U of catalase (Worthington Biochemical Corporation, New Jersey, USA) instead of 5% lysed horse blood. The lowest concentration of compound where no turbidity was observed was noted as the MIC. *S. pneumoniae* ATCC 49619 was used as quality control.

Minimum bactericidal concentration (MBC) of [Mn(CO)_3_(tpa-κ^3^*N*)]Br was determined using a resazurin-based microtiter plate assay as previously described (17). After adding 20 μL of 0.15 mg/mL resazurin (Cayman Chemical Company, Michigan, USA) solution in PBS to each well, plates were incubated at 37°C and the color conversion of all wells was recorded. The lowest well concentration of [Mn(CO)_3_(tpa-κ^3^*N*)]Br to remain blue was considered the MBC. All assays were performed in triplicate.

### Disc diffusion synergy test and checkerboard assays

Synergy between [Mn(CO)_3_(tpa-κ^3^*N*)]Br and 11 anti-pneumococcal agents was assessed by a modified disk diffusion test of the EUCAST method, in that a disk of each of the anti-pneumococcal agents was tested with and without the addition of 64 μg of [Mn(CO)_3_(tpa-κ^3^*N*)]Br. A decrease in the inhibition zone diameter for the combination discs versus the discs alone was considered suggestive of synergy.

To confirm the observed synergies and determine their magnitude, checkerboard assays were performed for three randomly selected strains (SP25, SP96 and SP30) using an inoculum of approximately 10^5^ CFU/ml onto each well and a 2-fold dilution scheme. The fractional inhibitory concentration (FIC) for each well and the FIC index were calculated as previously described (18). All assays were performed in triplicate.

### Time-kill assays

Time-kill assays were performed in triplicate using approximately 10^5^ CFU/mL as the starting inoculum for each strain and antimicrobials were added at the following final concentrations: [Mn(CO)_3_(tpa-κ^3^*N*)]Br (1 x MIC), tetracycline (1 x and 2 x MIC) and the Mn complex-tetracycline combination (0.5 x MIC – 1 x MIC). Cultures were incubated at 37°C under 5% CO_2_ continuous agitation (225 rpm) for 24 h. At set time points of 0, 30 min, 1, 2, 4 and 24 h post inoculation, 100 μL samples were collected, serially diluted and cultured onto blood agar plates for viable cell titer determination. Time-kill curves (CFU/ml vs time) were plotted using GraphPad Prism 8.2.1 software. Synergy was defined as bactericidal activity (≥2 log_10_ difference in CFU/mL) of the combination compared with either agent alone, after 24 h incubation. Unpaired student t-tests were performed to check for significant differences.

### *Galleria mellonella* treatment assays

*S. pneumoniae* inocula of approximately 0.3 OD_600_ (equating to ~10^8^ CFU/mL) in phosphate buffered saline (PBS) were serially diluted in PBS and colony forming units were determined by plating the dilutions on blood agar and incubating for 24 h. Sixteen *Galleria mellonella* larvae (TruLarv^™^, Biosystems Technology, Exeter, U.K) were infected with 10^5^ CFU/larvae of each *S. pneumoniae* strain (SP25, SP30 and SP96) via a 10 μL injection in a left proleg as previously described (19). Within 30 min of infection, a second injection into a right proleg was performed to administer the Mn complex (2.56 mg/kg in PBS), tetracycline (0.64 mg/kg), a combination of Mn complex and tetracycline (2.56 + 0.64 mg/kg) or PBS, respectively. Larvae were incubated at 37°C and scored for survival (live/dead) at 0, 24, 48, 72 and 96 h post inoculation.

Melanisation scores for larvae were recorded over 96 h as an indicator of morbidity, based on a reversed scoring method previously published (20), whereby a score of 4 indicated total melanisation of the larvae, 2 indicated melanin spots over the larvae, 1 indicated discoloration of the tail and a score of 0 indicated no melanisation.

All assays were performed in triplicate and the data was plotted using GraphPad Prism 8.2.1 software (San Diego, CA, USA). Analysis of survival curves was performed using the log rank test, with *a p* value of ≤ 0.05 indicating statistical significance (21). Unpaired student t-tests were performed to check for differences in bacterial counts at 24 h.

## Results and discussion

The antibacterial activity of [Mn(CO)_3_(tpa-κ^3^*N*)]Br was studied on 20 *S. pneumoniae* clinical isolates exhibiting genotypically-confirmed multidrug-resistance phenotypes (Table S1). [Mn(CO)_3_(tpa-κ^3^*N*)]Br was weak against *S. pneumoniae*, with MICs ranging from 64 mg/L (*n* = 11; 55%) to 128 mg/L (*n* = 9; 45%) (Table S1). However, this is 8-to 16-fold more active than previously shown against multidrug-resistant *E. coli* (14). This enhanced activity is potentially due to the absence of the Gram-negative outer lipopolysaccharide membrane known to reduce the permeability of many antimicrobials (22). The MBCs for all tested strains were equal to the MICs, suggesting bactericidal activity of [Mn(CO)_3_(tpa-κ^3^*N*)]Br. Time-kill assays for three randomly selected strains, SP25, SP30 and SP96, confirmed its bactericidal activity with total bacterial death observed within 2 h (SP25 and SP96) or 24 h (SP30) at 1x MIC (Figure 1a).

**Figure 1.**
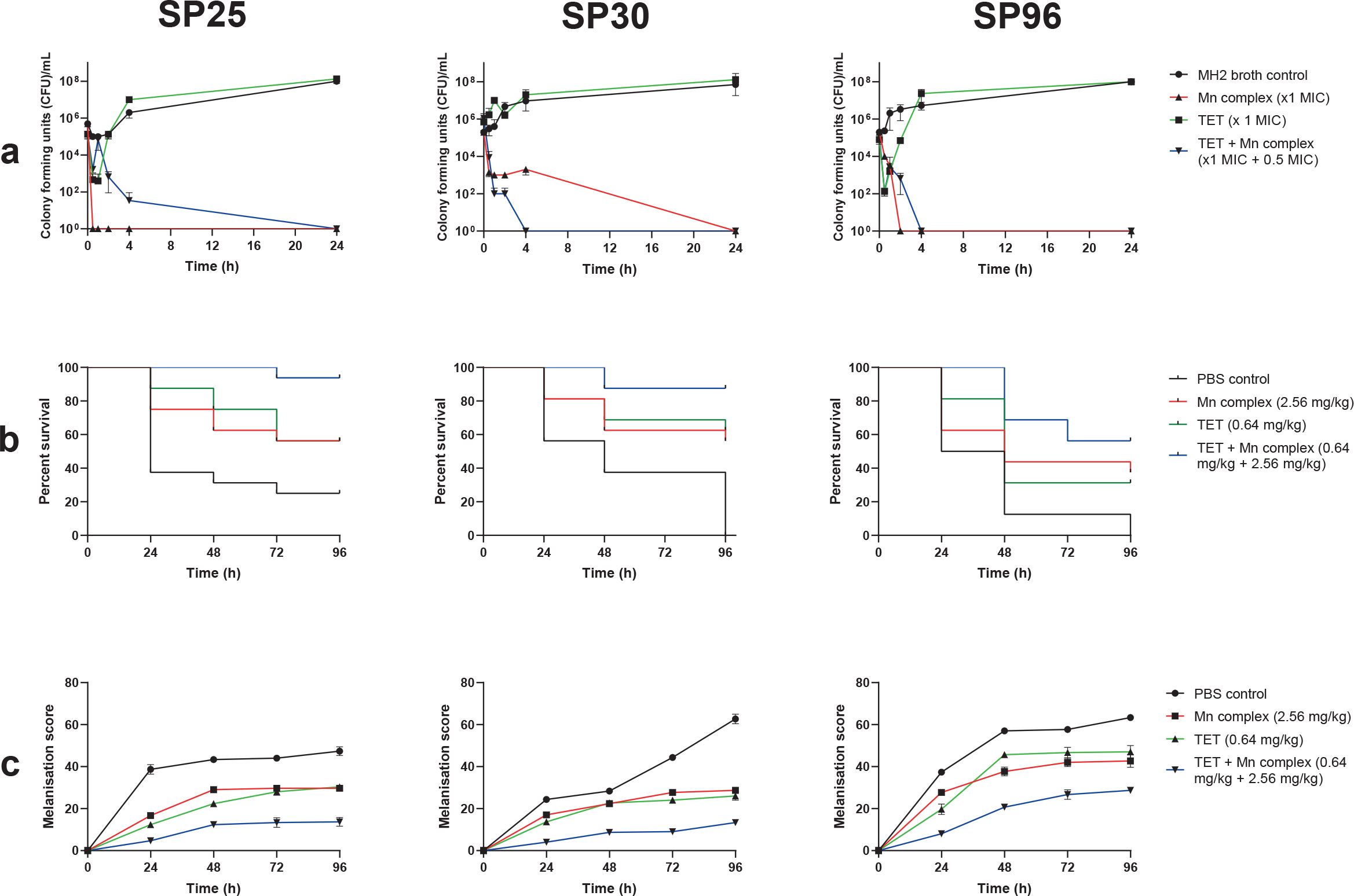
**(a)** Time-kill curves of [Mn(CO)_3_(tpa-κ^3^*N*)]Br, tetracycline and combination of both agents (x1 MIC + 0.5 MIC) versus *S. pneumoniae* strains SP25, SP30 and SP96 over 24 h. **(b)** Survival curves (live/dead) of *Galleria mellonella* over 96 h after infection with 10^5^ CFU/larvae of strains SP25, SP30 and SP96 and treatment with phosphate buffered saline (PBS), 2.56 mg/kg of [Mn(CO)_3_(tpa-κ^3^*N*)]Br, 0.64 mg/kg of tetracycline, and combination of both agents. **(c)** Melanisation assays in *G. mellonella* under the same conditions for strains SP25, SP30 and SP96.

The potential synergistic effect of the Mn complex with 11 other anti-pneumococcal agents against SP25, SP30 and SP96 was assessed by a combination disc diffusion test. All three strains showed a decreased diameter of inhibition zone only for the combinations of tetracycline, erythromycin and co-trimoxazole with the Mn complex versus these agents alone, suggesting synergy between these antibiotics and [Mn(CO)_3_(tpa-κ^3^*N*)]Br (Table S2). To examine strain-specific effects, we analyzed these same synergistic combinations for the remaining 17 multidrug-resistant *S. pneumoniae* strains by a combination disc diffusion test. Among them, eight (47.0%) exhibited decreased diameter of inhibition zone for tetracycline (ranging from 1 to 2 mm), 10 (58.8%) for erythromycin (ranging from 1 to 7.5 mm) and 13 (76.5%) for co-trimoxazole (ranging from 1 to 2 mm) (Table S2).

Checkerboard assays for SP25, SP30 and SP96 indicated that the Mn complex was able to increase susceptibility of tetracycline even against tetracycline-resistant strains of *S. pneumoniae*, with tetracycline MICs falling below the susceptibility breakpoint of 1 mg/L. Similar results showed that resistant strains were resensitized to erythromycin- and the co-trimoxazole-Mn complex combination (Table 1). Fractional inhibitory concentrate indexes were calculated and indicated that synergy was observed between co-trimoxazole and [Mn(CO)_3_(tpa-κ^3^*N*)]Br against all strains (FICI = 0.002-0.26) and against 2 out of 3 strains with combinations of [Mn(CO)_3_(tpa-κ^3^*N*)]Br with tetracycline (FICI = 0.123-0.28) and erythromycin (FICI = 0.28-0.31), with intermediate/additive activity observed with the remaining strains (0.75-2).

**Table 1.**
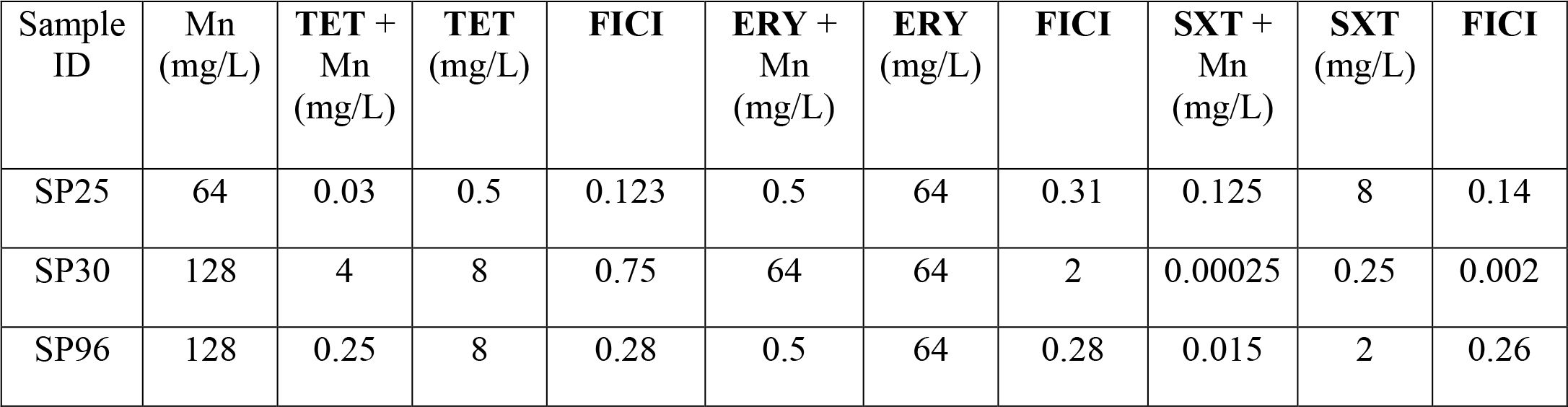
Minimum inhibitory concentrations (MICs) and fractional inhibitory concentration index (FICI) of the antibiotics tetracycline (TET), erythromycin (ERY) and co-trimoxazole (SXT) alone and in combination with the Mn complex [Mn(CO)_3_(tpa-κ^3^*N*)]Br against multidrug-resistant *S. pneumoniae* strains included in this study.

| Sample ID | Mn (mg/L) | TET + Mn (mg/L) | TET (mg/L) | FICI | ERY + Mn (mg/L) | ERY (mg/L) | FICI | SXT + Mn (mg/L) | SXT (mg/L) | FICI |
| --- | --- | --- | --- | --- | --- | --- | --- | --- | --- | --- |
| SP25 | 64 | 0.03 | 0.5 | 0.123 | 0.5 | 64 | 0.31 | 0.125 | 8 | 0.14 |
| SP30 | 128 | 4 | 8 | 0.75 | 64 | 64 | 2 | 0.00025 | 0.25 | 0.002 |
| SP96 | 128 | 0.25 | 8 | 0.28 | 0.5 | 64 | 0.28 | 0.015 | 2 | 0.26 |

Synergy between [Mn(CO)_3_(tpa-κ^3^*N*)]Br and tetracycline was confirmed using time-kill assays for the same strains, where a subinhibitory concentration of the Mn complex not only restored the activity of tetracycline, but was bactericidal. Bacteria were completely eradicated at 4 h (SP30 and SP96) and 24 h (SP25) with the combination, versus tetracycline alone, where an increase in bacterial numbers (10^7^ – 10^8^ CFU/mL) was observed at 24 h (Figure 1a). Previous studies have also highlighted synergy between doxycycline and [Mn(CO)_3_(tpa-κ^3^*N*)]Br against *E. coli* by reducing the expression of *tet(A)* (16). Therefore it is logical to postulate that in *tet(M)-encoding S. pneumoniae*, [Mn(CO)_3_(tpa-κ^3^*N*)]Br may reduce the expression of *tet(M)*, increasing susceptibility to tetracycline. Further studies are needed to confirm the mechanism of synergy in *S. pneumoniae*.

To evaluate the efficacy of the tetracycline-Mn complex combination *in vivo, G. mellonella* larvae were infected with *S. pneumoniae* strains SP25, SP30 and SP96. Overall, data from *in vivo* experiments show a significant difference between tetracycline monotherapy and the tetracycline-Mn complex combination *(p =* <0.049). With only PBS therapy, infections with SP25, SP30 and SP96 resulted in mortality rates of 75%, 62.5% and 87.5%, respectively (Fig 1b), reflecting intrinsic differences in strain virulence. Treatment with the tetracycline-Mn complex combination was superior to monotherapy with either tetracycline or the Mn complex, resulting in significantly lower mortality in infected larvae (20.8% vs 47.9% and 45.8% respectively). Consistent with mortality data, high melanisation scores indicated a strong immune response in *G. mellonella* infected with strains SP25, SP30 and SP96 (Fig. 1c), with mean scores of 47 (+/- 2.3), 63 (+/- 2.7) and 63 (+/- 1.3) out of a maximum of 64 for each strain, respectively. Melanisation was reduced in larvae treated with the tetracycline-Mn complex combination, compared with tetracycline and Mn complex monotherapy, with mean scores of 18.6 (+/- 7.2), 34.4 (+/- 9.1) and 33.7 (+/- 6.4) respectively. Doses of [Mn(CO)_3_(tpa-κ^3^*N*)]Br used in this study for treatment of *S. pneumoniae* infections, were more than 70 times lower than the concentration previously shown to be toxic 24 h post-administration in *G. mellonella* (14).

In conclusion, our results show that [Mn(CO)_3_(tpa-κ^3^*A*)]Br used in combination with traditional antibiotics like tetracycline, erythromycin and co-trimoxazole, may have potential as antimicrobial and resistance breaker against multidrug-resistant *S. pneumoniae*.

## Acknowledgements

The authors are grateful to Dr. Lesley McGee and CDC for providing us with the clinical strains included in the study.

## Transparency declarations

None to declare.

## References

1. Lynch JP, Zhanel GG. 2009. *Streptococcus pneumoniae:* Epidemiology, Risk Factors, and Strategies for Prevention. Seminars in Respiratory and Critical Care Medicine 30:189–209.

2. Black RE, Cousens S, Johnson HL, Lawn JE, Rudan I, Bassani DG, Jha P, Campbell H, Walker CF, Cibulskis R, Eisele T, Liu L, Mathers C, Unicef W. 2010. Global, regional, and national causes of child mortality in 2008: a systematic analysis. Lancet 375:1969–1987.

3. World Health Organization. 2014. Global immunization data. World Health Organization, Geneva, Switzerland.

4. Pelton SI, Huot H, Finkelstein AA, Bishop CJ, Hsu KK, Kellenberg J, Huang SS, Goldstein R, Hanage WP. 2007. Emergence of 19A as virulent and multidrug resistant pneumococcus in Massachusetts following universal immunization of infants with pneumococcal conjugate vaccine. Pediatric Infectious Disease Journal 26:468–472.

5. Kim L, Mcgee L, Tomczyk S, Beall B. 2016. Biological and Epidemiological Features of Antibiotic-Resistant *Streptococcus pneumoniae* in Pre- and Post-Conjugate Vaccine Eras: a United States Perspective. Clinical Microbiology Reviews 29:525–552.

6. Lalitha MK, Pai R, Manoharan A, Appelbaum PC, Group CPS. 2002. Multidrug-resistant *Streptococcus pneumoniae* from India. Lancet 359:445.

7. Mera RM, Miller LA, Daniels JJ, Weil JG, White AR. 2005. Increasing prevalence of multidrug-resistant *Streptococcus pneumoniae* in the United States over a 10-year period: Alexander Project. Diagn Microbiol Infect Dis 51:195–200.

8. Overweg K, Hermans PWM, Trzcinski K, Sluijter M, De Groot R, Hryniewicz W. 1999. Multidrug-resistant *Streptococcus pneumoniae* in Poland: Identification of emerging clones. Journal of Clinical Microbiology 37:1739–1745.

9. Rahman M, Hussain S, Shorna S, Rashid H, Baqui AH, van der Linden M, Al-Lahham A, Reinert RR. 2006. Emergence of a unique multiply-antibiotic-resistant *Streptococcus pneumoniae* serotype 7B clone in Dhaka, Bangladesh. Journal of Clinical Microbiology 44:4625–4627.

10. Thummeepak R, Leerach N, Kunthalert D, Tangchaisuriya U, Thanwisai A, Sitthisak S. 2015. High prevalence of multi-drug resistant *Streptococcus pneumoniae* among healthy children in Thailand. Journal of Infection and Public Health 8:274–281.

11. Wang CY, Chen YH, Fang C, Zhou MM, Xu HM, Jing CM, Deng HL, Cai HJ, Jia K, Han SZ, Yu H, Wang AM, Yin DD, Wang CQ, Wang W, Huang WC, Deng JK, Zhao RZ, Chen YP, Yang JH, Wang C, Che YR, Nie XZ, Wang SF, Hao JH, Zhang CH. 2019. Antibiotic resistance profiles and multidrug resistance patterns of *Streptococcus pneumoniae* in pediatrics A multicenter retrospective study in mainland China. Medicine 98.

12. Guntzel P, Nagel C, Weigelt J, Betts JW, Pattrick CA, Southam HM, La Ragione RM, Poole RK, Schatzschneider U. 2019. Biological activity of manganese(i) tricarbonyl complexes on multidrug-resistant Gram-negative bacteria: From functional studies to *in vivo* activity in *Galleria mellonella*. Metallomics 11:2033–2042.

13. Nagel C, McLean S, Poole RK, Braunschweig H, Kramer T, Schatzschneider U. 2014. Introducing [Mn(CO)_3_(tpa-kappa^3^*N*)](+) as a novel photoactivatable CO-releasing molecule with well-defined iCORM intermediates - synthesis, spectroscopy, and antibacterial activity. Dalton Trans 43:9986–97.

14. Betts J, Nagel C, Schatzschneider U, Poole R, La Ragione RM. 2017. Antimicrobial activity of carbon monoxide-releasing molecule [Mn(CO)_3_(tpa-kappa^3^*N*)]Br versus multidrug-resistant isolates of Avian Pathogenic *Escherichia coli* and its synergy with colistin. PLoS One 12:e0186359.

15. Tinajero-Trejo M, Rana N, Nagel C, Jesse HE, Smith TW, Wareham LK, Hippler M, Schatzschneider U, Poole RK. 2016. Antimicrobial Activity of the Manganese Photoactivated Carbon Monoxide-Releasing Molecule [Mn(CO)_3_(tpa-kappa^3^*N*)](+) Against a Pathogenic *Escherichia coli* that Causes Urinary Infections. Antioxidants & Redox Signaling 24:765–780.

16. Rana N, Jesse HE, Tinajero-Trejo M, Butler JA, Tarlit JD, zur Muhlen MLU, Nagel C, Schatzschneider U, Poole RK. 2017. A manganese photosensitive tricarbonyl molecule [Mn(CO)_3_(tpa-kappa^3^*N*)]Br enhances antibiotic efficacy in a multi-drug-resistant *Escherichia coli*. Microbiology-Sgm 163:1477–1489.

17. Riss TL, Moravec RA, Niles AL, Duellman S, Benink HA, Worzella TJ, Minor L. 2004. Cell Viability Assays. In Sittampalam GS, Grossman A, Brimacombe K, Arkin M, Auld D, Austin CP, Baell J, Bejcek B, Caaveiro JMM, Chung TDY, Coussens NP, Dahlin JL, Devanaryan V, Foley TL, Glicksman M, Hall MD, Haas JV, Hoare SRJ, Inglese J, Iversen PW, Kahl SD, Kales SC, Kirshner S, Lal-Nag M, Li Z, McGee J, McManus O, Riss T, Saradjian P, Trask OJ, Jr., Weidner JR, Wildey MJ, Xia M, Xu X (ed), Assay Guidance Manual, Bethesda (MD).

18. Hall MJ, Middleton RF, Westmacott D. 1983. The Fractional Inhibitory Concentration (Fic) Index as a Measure of Synergy. Journal of Antimicrobial Chemotherapy 11:427–433.

19. Evans BA, Rozen DE. 2012. A *Streptococcus pneumoniae* infection model in larvae of the wax moth *Galleria mellonella*. Eur J Clin Microbiol Infect Dis 31:2653–60.

20. Betts JW, Hornsey M, Wareham DW, La Ragione RM. 2017. *In vitro* and *In vivo* Activity of Theaflavin-Epicatechin Combinations versus Multidrug-Resistant *Acinetobacter baumannii*. Infectious Diseases and Therapy 6:435–442.

21. Hornsey M, Wareham DW. 2011. *In vivo* efficacy of glycopeptide-colistin combination therapies in a *Galleria mellonella* model of *Acinetobacter baumannii* infection. Antimicrob Agents Chemother 55:3534–7.

22. Delcour AH. 2009. Outer membrane permeability and antibiotic resistance. Biochim Biophys Acta 1794:808–16.

